# Not like other conifers: evaluation of phenotypic diversity in British common juniper, *Juniperus communis*, indicates genetic isolation and local adaptations among remnant populations

**DOI:** 10.1101/2024.09.26.615138

**Authors:** J. Baker, J. Cottrell, R. Ennos, A. Perry, S. Green, S. Cavers

## Abstract

Habitat fragmentation and genetic isolation pose threats to the genetic diversity and resilience of natural populations. Protecting the genetic diversity of populations, and the processes that sustain it, optimises their ability to adapt to changing conditions and new threats: conservation efforts with this specific goal are known as “dynamic conservation.” The common juniper, *Juniperus communis*, is a keystone species that provides habitat and resources for many plants and animals. It is a highly polymorphic species, and across its natural range it grows in a variety of habitats and growth forms. Juniper populations have been shrinking and becoming increasingly fragmented for over a century in the UK and elsewhere in Europe, raising concerns about the genetic diversity present in juniper populations and their ability to adapt to changing conditions, or their adaptive potential. This paper presents an analysis of the partitioning of phenotypic diversity among regions, populations and families from 16 UK populations assessed in a common garden trial. Our findings suggest high phenotypic variation among populations compared to the variation among families within populations, indicating barriers to gene flow between juniper populations, relatively homogenous populations and, consequently, potentially reduced adaptive potential. This information is a useful baseline for conservation managers and can also help to infer the genetic diversity and adaptive potential of populations.

## Introduction

**Figure 1:**
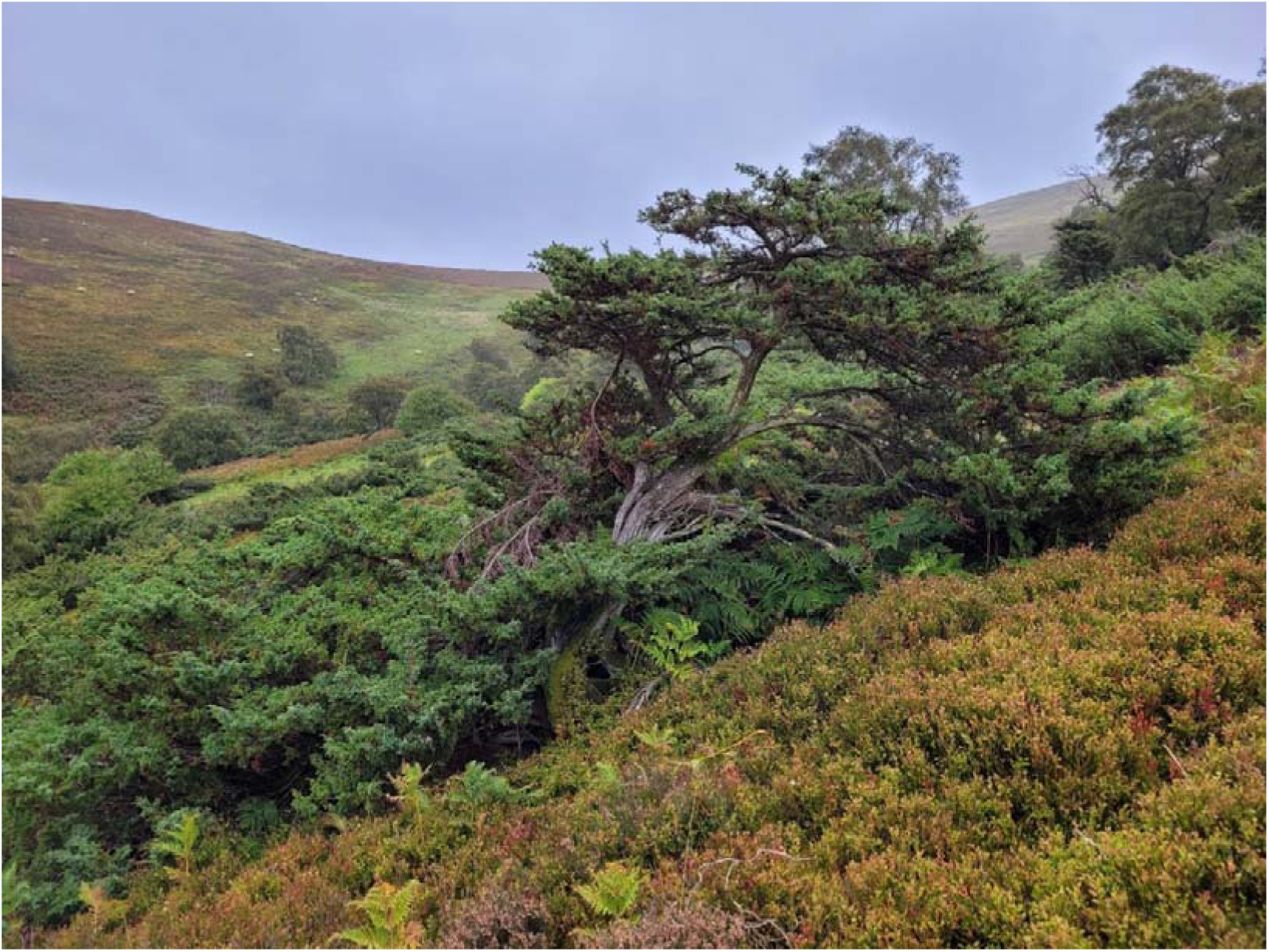
cover photo taken by Baker, **J.**

The capacity of populations to adapt to changing conditions is underpinned by genetic variation and the extent to which it is heritable across generations (Boyd et al., 2022; Geng et al., 2016). Maintaining genetic diversity by ensuring a dynamic balance between gene flow and natural selective processes can support the adaptive potential and therefore resilience of populations and species. Strategies that aim to do so are typically termed “dynamic conservation” (Cavers & Cottrell, 2015; Eriksson et al., 1993), and they have been adopted as a genetic resource conservation approach, particularly for tree species (Lefèvre et al., 2013). Here, we aim to evaluate the adaptive potential of the UK populations of common juniper, *Juniperus communis,* using a common garden trial. Juniper is one of only three conifers native to the UK, and despite intensive conservation efforts in some areas such as in Southern England, many populations are shrinking and becoming increasingly fragmented. Some populations have declined to the point of senescence and have failed to reproduce naturally in recent years. This population decline may have serious repercussions for the genetic diversity, and therefore adaptive potential and resilience, of the UK’s remaining juniper populations, especially given the changing climate and the introduction of non-native pathogens. We quantified phenotypic diversity in 16 juniper populations to gain an understanding of the standing genetic variation of populations, or the amount of “raw [genetic] material” (Aguilar et al., 2008) upon which natural selection can act. As adaptive potential depends on phenotypic variation (changing conditions will impose selection, and selection acts on the phenotype), the most valuable assessment of genetic diversity is to quantify the genetic component of phenotypic variation. This information will help conservation managers understand how juniper may tolerate both its current and future stressors.

Common garden trials are used by researchers to assess the genetic variation underpinning phenotypic traits (de Villemereuil et al., 2016; Ramírez□Valiente et al., 2022; Schwinning et al., 2022). Since a phenotype is a product of both genes and environment, common garden trials allow quantification of the genetic part of phenotypic trait variation by minimizing the effects of the environment. This information can be used to identify locally adaptive traits (McKay et al., 2001) and guide conservation efforts, including informing the debate about the need to transplant individuals between populations in such a way as to minimize the risks of maladaptation (Schwinning et al., 2022). Furthermore, if the trial design incorporates pedigree information (such as family identity), heritability can be estimated to elucidate the patterns of local adaptation that selection may act upon (de Villemereuil et al., 2016; Donnelly et al., 2016; Schwinning et al., 2022). Finally, patterns of variation in putatively adaptive traits identified in common garden trials can be correlated with local climate variables from the source populations to improve our understanding of the climatic drivers of local adaptation (López et al., 2008; McKay et al., 2001; RamírezlJValiente et al., 2022; Schwinning et al., 2022). This paper evaluates genetically based phenotypic diversity both within and among progeny from 16 natural UK juniper populations using a common garden trial to provide a genetic baseline assessment of standing phenotypic genetic diversity in this UK Biodiversity Action Plan priority species.

The common juniper is the most widespread conifer in the world, with a circumpolar range spanning from the tundras of Russia and Canada as far south as the Mediterranean in Europe and the Central Rocky Mountains in North America (Thomas et al., 2007). Across this range, juniper displays striking phenotypic variability that has been hypothesized to be a result of both phenotypic plasticity as well as distinct genetic adaptations (Knyazeva & Hantemirova, 2020; Sullivan, 2001; Thomas et al., 2007; Ward, 2007). For example, *J. communis* can grow as upright mid-story trees, sprawling shrubs or prostrate ground-hugging stems depending on its environment (Sullivan, 2001; Thomas et al., 2007). Some researchers have hypothesized that it is juniper’s phenotypic plasticity that enables it to survive in such a broad suite of habitats and act as a pioneer species after natural disturbances (Knyazeva & Hantemirova, 2020; Thomas et al., 2007). The degree to which the observed phenotypic variation within *J. communis* is due to environmental and/or genetic differences is unclear.

Three subspecies of the common juniper occur in the UK: *J. communis* ssp. *communis*, *J. communis* ssp. *nana*, and *J. communis* ssp. *hemisphaerica*. *J. communis* ssp. *communis* has the widest range of the three subspecies in the UK, and populations of ssp. *nana* and ssp. *hemisphaerica* are restricted to the west coast of Scotland and the Lizard Peninsula in Southern England, respectively. Although the genetic status of these three subspecies is unclear, the phenotypic and genetic diversity of Scottish *J. communis* and *J. communis* ssp. *nana* populations were investigated by Sullivan (2001). They found that *J. communis* ssp. *communis* was capable of phenotypic plasticity, with prostrate cuttings from *J. communis* ssp. *communis* plants from harsh environments growing as shrubs or upright trees in the greenhouse, whereas *J. communis* ssp. *nana* did not exhibit this plasticity and maintained its prostrate habit in the greenhouse. Although they did not find clear evidence of a genetic distinction between ssp. *communis* and ssp. *nana,* they did find evidence that both the growth habit of ssp. *nana* is a genetic adaptation as well as a notably plastic response of ssp. *communis* to its environment. Here, we include one population (AR) that would be considered to be ssp. *nana* based on the work of Sullivan (2001), and our work builds on this study by: including more than twice as many populations; extending the geographic range of populations beyond Scotland to include populations from both the Lake District and Southern England; and by using information on the pedigrees of individuals to elucidate the degree of genetically-based phenotypic variance in *J. communis* using a common garden trial.

Despite the fact that juniper’s phenotypic variation would seem to suggest a highly adaptive and resilient species, juniper populations across the UK and parts of Europe have been in a state of decline for at least the past century (Sullivan, 2003; Verheyen et al., 2009). Juniper occurs across the UK from Cornwall to Shetland but has substantial populations in three main geographic regions: Southern England, the Lake District, and Scotland. Southern English populations are particularly small and fragmented, whereas populations in the Lake District and Scotland tend to be larger. Many populations across the UK are suffering reproductive failure (Sullivan, 2001; Thomas et al., 2007). Understanding the genetic diversity within these different regions and populations is especially important as a basis for dynamic conservation, and to anticipate and mitigate the impacts of habitat fragmentation and reproductive senescence.

Here, we quantified patterns of within and between family variation in phenotypic traits of *J. communis* from populations across Britain using a common garden trial. We also compared variation between the three main British population centres, or regions, which were identified as genetic units in a parallel analysis of the population genetics of common junipers in Britain (Baker *et. al*, 2024; in preparation). Finally, we compared the phenotypic data from our common garden trial with climate data from source population locations. The main goals of this work were:

1. to estimate the extent and pattern of quantitative genetic trait variation, particularly how variation is partitioned between regions, populations and families;
2. to identify the primary traits showing among-population variation, suggesting that they may be important for local adaptation;
3. to identify the key abiotic factors driving local adaptation by correlating quantitative trait variation with climate variables from the source population’s location.

## Methods

### Seed collection

Seeds were collected in autumn 2015 from natural populations in England and Scotland (Table 1, Figure 2). Although the UK distribution of juniper includes populations in Wales and Ireland, no samples from these areas were included in this study. Both population and family identities were recorded for each seedlot. Population refers to the stand of juniper from which the berries were collected, and family refers to a collection of open-pollinated seed from one maternal tree, most likely half-siblings. Berries were collected from a minimum of four and an average of eight well-spaced maternal trees from each population. Berries were stored at 4°C until they were ready to be processed and seeds were extracted from berries by hand. Seed viability was evaluated using the float test, where seeds that sank in water were considered viable and those that floated were assumed to be empty and therefore nonviable. A total of 26,585 viable seeds were sown (full data are available from the EIDC archive (Baker et al., 2024a).

**Figure 2:**
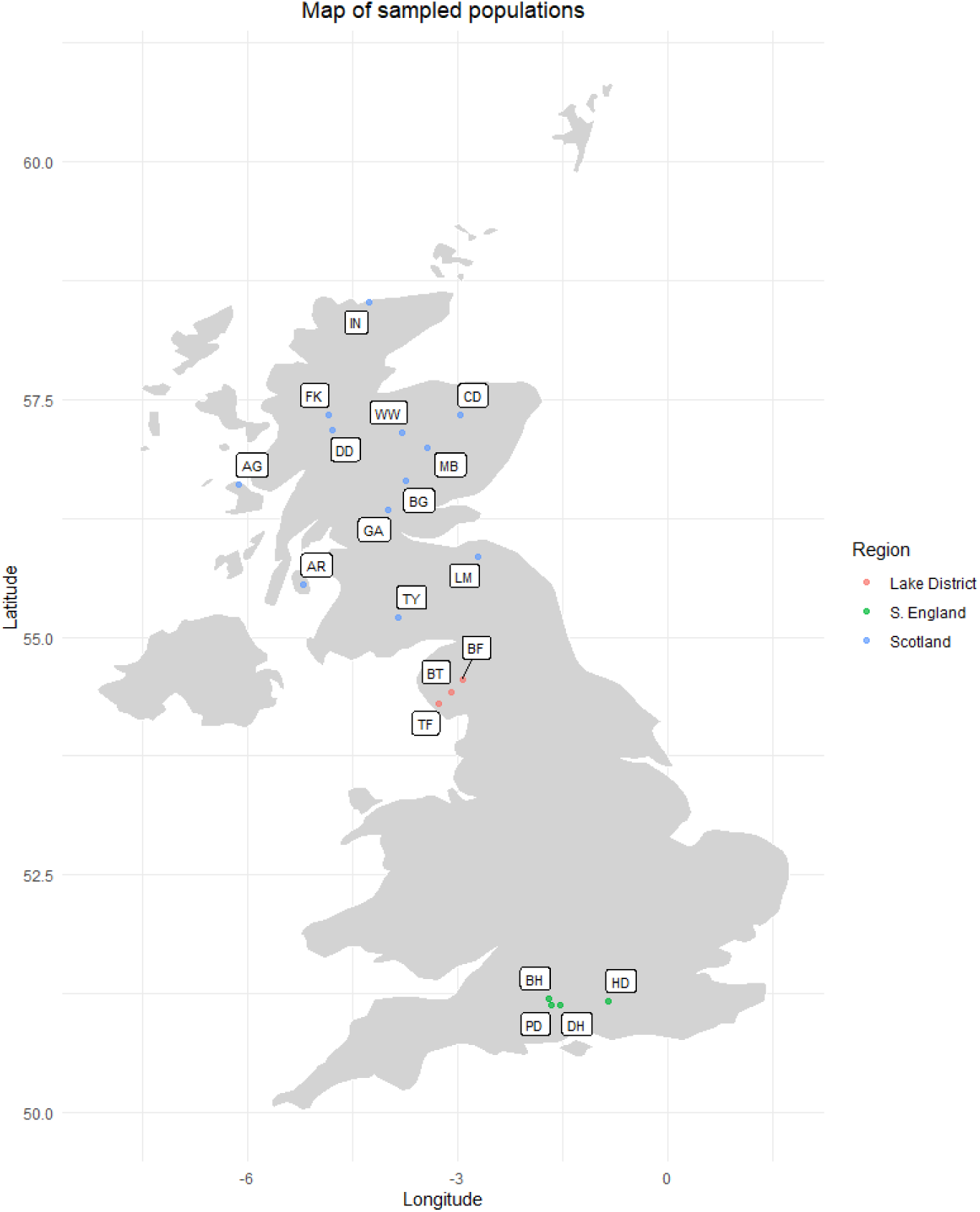
Map displaying locations of populations from which seeds were collected for common garden trial. Colours denote: red-Scotland, green-Lake District, blue-S. England

**Table 1:** List of populations collected for inclusion in the greenhouse trial with abbreviations, locations, the number of families from which seeds were collected and the number of individuals that were ultimately included in the common garden trial.

| Population | Abbreviation | Region | Latitude (°) | Longitude (°) | Number of families | Number of individuals |
| --- | --- | --- | --- | --- | --- | --- |
| Argyll | AG | Scotland | 56.605 | -6.123 | 7 | 20 |
| Gleann Dubh, Arran | AR | Scotland | 55.550 | -5.202 | 7 | 21 |
| Blowick Fell | BF | Lake District | 54.558 | -2.931 | 9 | 30 |
| Balnaguard Glen | BG | Scotland | 56.644 | -3.730 | 8 | 27 |
| Bulford Hill | BH | S. England | 51.204 | -1.700 | 2 | 6 |
| Blea Tarn | BT | Lake District | 54.426 | -3.088 | 8 | 24 |
| Clashindarroch | CD | Scotland | 57.337 | -2.966 | 10 | 0 |
| Dundreggan | DD | Scotland | 57.185 | -4.790 | 2 | 6 |
| Danebury Hill | DH | S. England | 51.137 | -1.534 | 10 | 0 |
| Fasnakyle | FK | Scotland | 57.336 | -4.849 | 1 | 3 |
| Glenartney | GA | Scotland | 56.340 | -3.996 | 7 | 27 |
| Harting Down | HD | S. England | 51.178 | -0.851 | 4 | 0 |
| Invernaver | IN | Scotland | 58.521 | -4.256 | 3 | 9 |
| Lammermuir | LM | Scotland | 55.853 | -2.710 | 9 | 26 |
| Morrone Birkwood | MB | Scotland | 56.998 | -3.426 | 5 | 15 |
| Porton Down | PD | S. England | 51.138 | -1.652 | 1 | 3 |
| Thwaites Fell | TF | Lake District | 54.302 | -3.263 | 11 | 39 |
| Tynron | TY | Scotland | 55.214 | -3.846 | 2 | 6 |
| Whitewell | WW | Scotland | 57.155 | -3.796 | 7 | 27 |

### Seed stratification

Seeds were initially sown in seed trays (6cm x 24cm x 38cm (H x W x L) with drainage holes) using a 1:1 sand:compost soil mix in December 2015. Trays were placed in loose plastic bags and subjected to a warm/ cool/ warm stratification protocol (MacCartan, 2013) in the greenhouse at the UKCEH, Edinburgh (latitude 55.86°, longitude -3.21°). The stratification protocol comprised of: ambient light at 7.3-14.3°C from December 2015-March 2016 followed by a cold, dark stratification period at 4°C from March 2016-October 2016, and finally, the seed trays were placed at ambient temperature and light in a greenhouse. Seedlings emerged sporadically from October 2016 to August 2018, after which the seed trays were disposed of. During four potting up days (October 2016, February 2017, October 2017 and August 2018), any seedlings that had emerged were picked out and transplanted into 8×8×8 cm pots with a 1:1 sand: compost mix and placed on an irrigated bench in the greenhouse at ambient light and temperature. Therefore, there were four discrete age classes, hereafter denoted 1-4, with one being the first (oldest) and four being the last (youngest) plants to emerge. These age classes were recorded for each individual to allow for the variation in traits caused by plant age to be accounted for in subsequent analyses. Each plant was labelled with its population, family, individual identity and age class.

### Common garden experiment

The common garden experiment consisted of 89 families from 16 English and Scottish populations in a randomised complete block design with one replicate in each of three blocks that was set up in the greenhouse. In December 2018 the 89 families that had at least three surviving individuals were re-potted into 13×13×13 cm pots. The populations CD, DH and HD were excluded at this point because no families from these populations had 3 surviving individuals. Up to 12 plants from each population were re-potted. Six families whose seedlings emerged in two different potting waves were represented by three individuals in each of multiple age classes: they were BF103 (age classes 1 and 4), BG1 (age classes 2 and 4), GA123 (age classes 1, 2 and 4), TF13 (age classes 1 and 4), TF5 (age classes 1 and 4) and WW2 (age classes 1, 2 and 4). All other families were represented by one individual per block, meaning that the first three blocks were each composed of 97 plants, and the remaining 9 blocks consisted of between 56-82 individuals, depending on seedling availability.

Traits were measured on plants from the three complete blocks, which had representatives from all 89 families. Two families, AG7 and LM14, were only represented by two individuals due to plants dying after being re-potted in December 2018, but are nevertheless included in these analyses. Populations were represented by between one (FK and PD) and 11 (TF) families each based on seedling availability (Table 1). Therefore, traits were measured on 289 plants from 89 families representing 16 populations.

Eleven putatively adaptive traits that were identified by Sullivan (2001) were measured during December 2020. They are described in Table 2. Block and age class were also recorded for each plant to include as control factors in the subsequent analyses. All traits were measured in the three complete blocks with the exception of needle length and needle width, which were not recorded in block 1. Seven and nine families were removed from the analyses of needle length and width, respectively, due to only having a measurement from one individual. The removed families were: AG7, BF106, BH9, IN3, LM22, MB8 and TY6 for needle length and AG1, AG7, BF106, GA128, IN3, LM22, MB8 and TY6 for needle width.

**Table 2:**
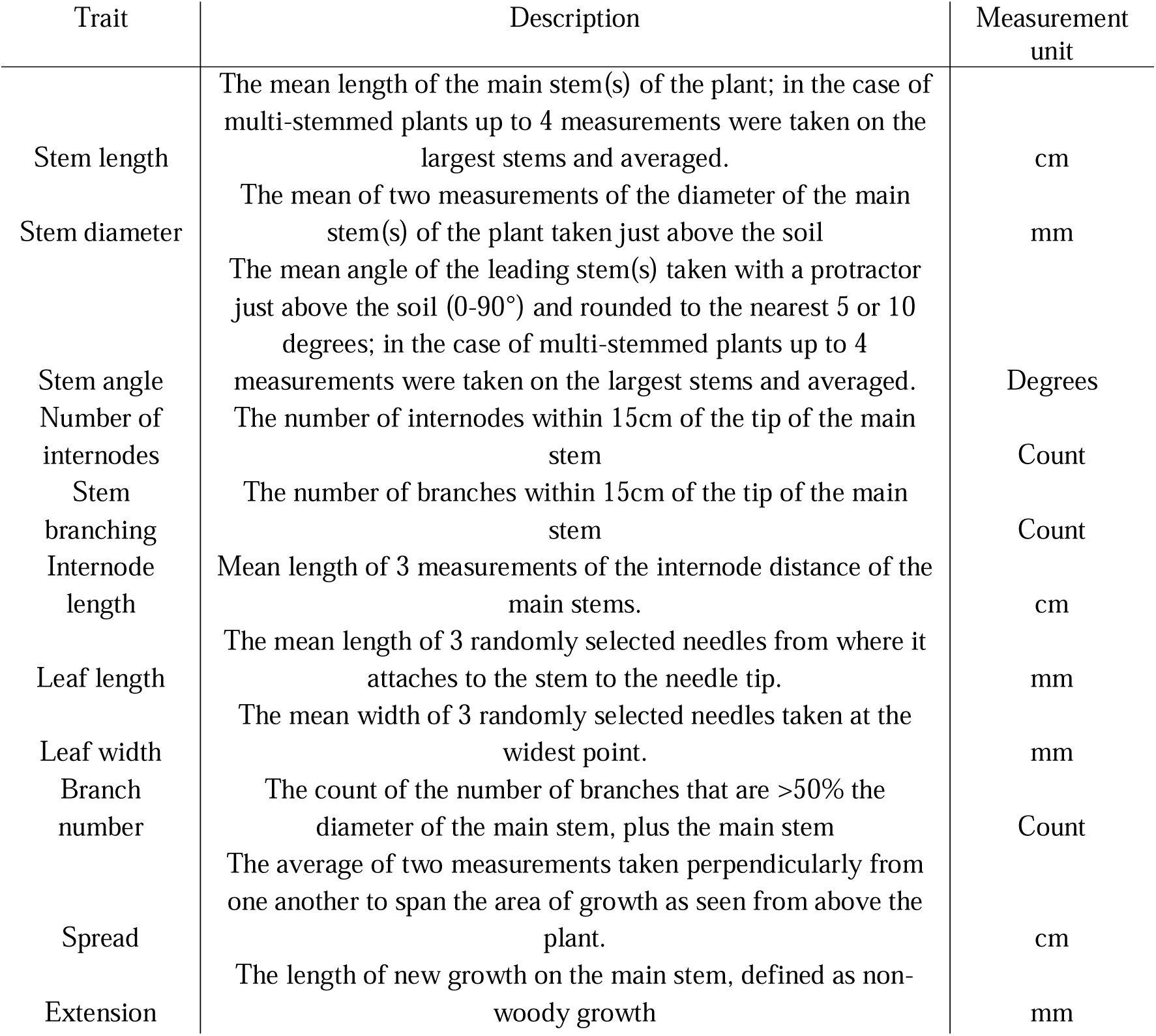
Summary of all 11 measured traits, including brief descriptions of how traits were measured.

### Analyses of trait data

Statistical analyses were conducted using Minitab 2019 (*Minitab 19*, 2024). Trait data were checked for outliers and skewedness; extreme outliers were removed before running Grubbs outlier tests and Shapiro-Wilks normality tests on all traits. Branch number and spread had additional outliers that were removed, and branch number was square-root transformed to correct for right skewness. To test for variation between and within regions and populations, general linear models were run using region as a fixed effect, population as a fixed effect nested within region, family as a random effect nested within population, age class as a fixed effect and block as a random effect. To test for variation within age classes, general linear models were run within the youngest and oldest age classes using the same factors and nesting as above, excluding age class. A Bonferroni correction for multiple tests was applied to all general linear models, giving a revised significance level of p=0.0045. The results of these models can be found in Appendix 1.

Proportional Expected Means Squared (PEMS) values were calculated for each predictor except the control factors (block and age class) from Adjusted Means Squared (AMS) values by subtracting the AMS of each predictor from the level of nesting “below” it and dividing that by the average number of individuals in that nesting level. For example, PEMS of families is equal to the AMS for families, minus the AMS for the error term divided by the average number of individuals per family. The sum of these values was used to calculate the PEMS for each predictor (Figure 3).

**Figure 3:**
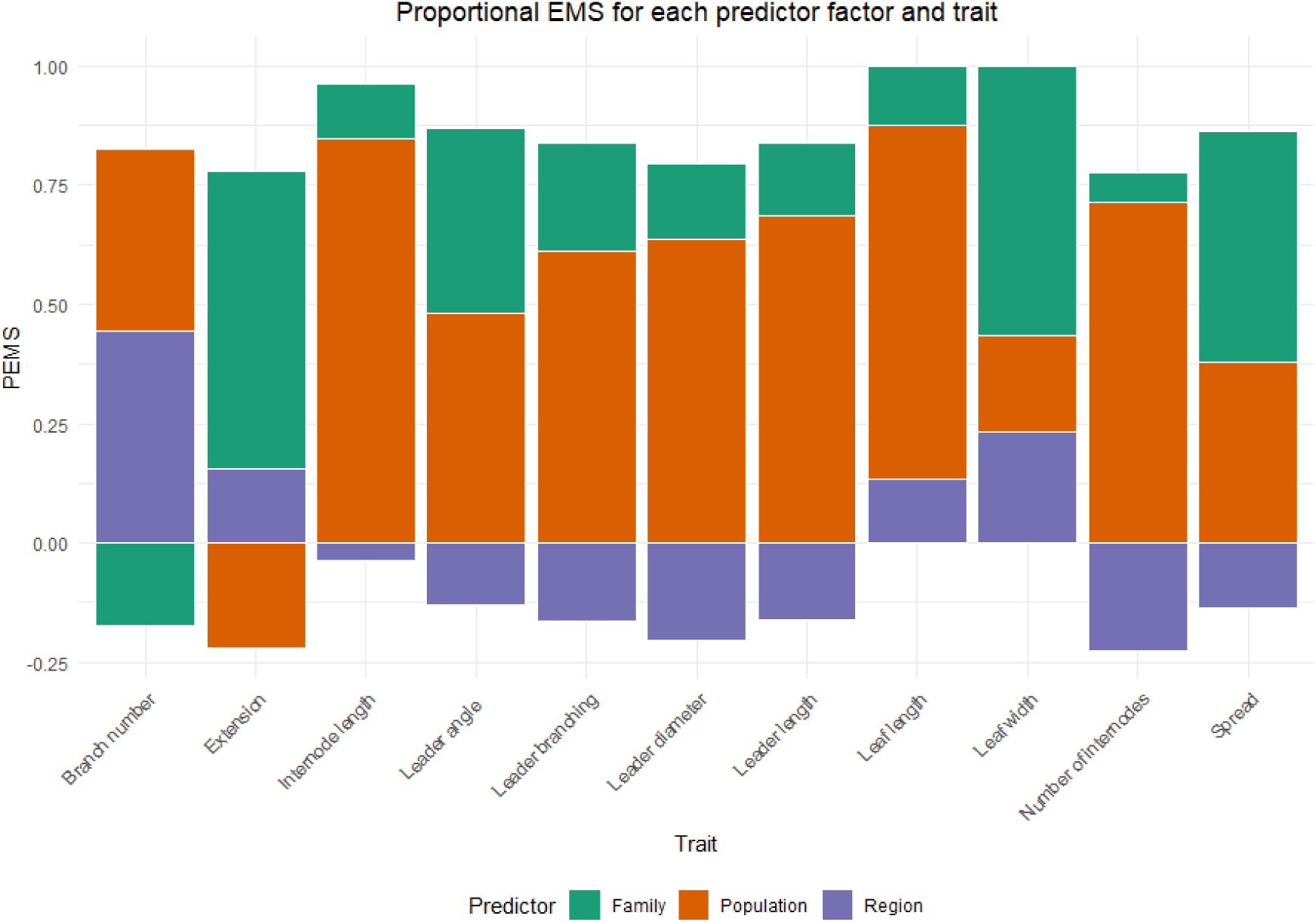
Proportional Expected Means Squared (PEMS) values for region, population and family for each trait, presented as percentages of the sum of expected means squared values. Represents the proportion of the observed variation that each predictor factor was ascribed by general linear models. A negative PEMS value indicates that inclusion of the predictor did not improve the model.

Narrow-sense heritability (*h^2^*) was calculated by multiplying the EMS of family by four to account for average sibling relatedness and dividing by the sum of EMS values for all predictors and the error term. For example, the *h^2^* of stem length is equal to (EMS Family*4)/Sum EMS= (9.7*4)/145.9=0.265.

### Accessing and processing climate data

Correlations between trait measurements and climate variables were conducted only for Scottish populations as this removed the potentially confounding effect of regional population genetic structure (Baker *et. al*, 2024; in preparation) while maintaining sample size. Data on climate variables for the 30-year period 1989-2019 were gathered from HadUK-Grid (Hollis et al., 2018) on a 1km^2^ resolution for each of the 9 Scottish populations. We included: average February minimum temperature, average annual cumulative rainfall, average annual temperature and average annual cumulative days with ground frost. Monthly means for each variable were converted from .nc to .csv files using the “raster” package for R (Hijmans, 2023), then processed into annual means as needed on base R (Team, 2021). Processed values that were used in the proceeding Principal Component Regressions (PCR) for each population can be found in Appendix 2.

### Regression of trait data on climate variables

Principal Component Regressions (PCRs) were used to evaluate the influence of climate at the site origin on trait variations. PCR is capable of handling multicollinearity among predictor variables (Kovoor & Nandagiri, 2007) and is commonly used to simplify complex climatic datasets and identify the most important predictors (Marenco & Antezana-Vera, 2021). Minitab 19 (*Minitab 19*, 2024) was used to run a Principal Component Analysis (PCA) on the four climate variables. The first two principal components were used as predictor variables in regression analyses with each of the 11 traits. Since climate data were on a 1km^2^ resolution, which typically encompasses the entire areas of the source populations, trait data were averaged by population to compare to the climate data. To control for differences between age classes, these regressions were run within both the youngest and oldest age class. To account for the number of tests that were performed, a Bonferroni correction was applied, giving a revised significance level of p=0.0045.

## Results

### Variation of trait data

The main contributor to the observed variation in six of the eleven traits was the difference between populations (Figure 3). Although region was only a significant predictor for branch number, the inclusion of regions improved the models for leaf length, leaf width, branch number and extension. The six traits for which population was significant were: stem length (p=3.70E-07), stem diameter (p=1.05E-05), number of internodes (p=1.45E-04), stem branching (p=1.67E-05), internode length (p=2.87E-04) and leaf length (p=3.90E-05). Finally, family was significant for leaf width (p=3.37E-03) and spread (p=2.76E-03) and accounted for a large portion of the variation in both. Both age class and block were important control factors, being significant predictors for four and five traits, respectively, and so both were retained in all models (Table 3).

**Table 3:** Results of general linear models and calculated narrow-sense heritability (H2) comparing 11 phenotypic traits measured from a mmon garden trial of 289 *J. communis* plants. Tests were nested general linear models using regions (fixed factor), populations (fixed factor) nested within regions, families (random factor) nested within populations, age class as a fixed control factor and block as a random control factor. Stars next to adjusted MS values indicate significance levels: key: * = p-value between 0.0045 and 0.00045, ** = p-value between 0.00045 and 0.000045, *** = p-value <0.000045.

|  |  | Stem length | Stem diameter | Stem angle | Number of internodes | Stem branching | Internode length | Leaf length | Leaf width | Branch number | Spread | Extension |
| --- | --- | --- | --- | --- | --- | --- | --- | --- | --- | --- | --- | --- |
| Model adjusted R<sup>2</sup> |  | 45.40% | 45.54% | 11.94% | 16.87% | 24.11% | 24.58% | 36.9% | 47.5% | 5.40% | 44.94% | 15.1% |
| Region | Adjusted MS | 87.72 | 0.7927 | 0.420368271 | 28.09 | 30.077 | 0.0383536 | 21.169 ** | 0.18484 ** | 0.492 | 75.48 | 1202.7 |
|  | DF | 2 | 2 | 2 | 2 | 2 | 2 | 2 | 2 | 2 | 2 | 2 |
| Population | Adjusted MS | 742.21*** | 14.6362** | 949.3 | 107.77** | 129.971*** | 25.8701* | 9.53** | 0.07015 | 0.2272 | 231.29 | 873.4 |
|  | DF | 11 | 11 | 11 | 11 | 11 | 11 | 11 | 11 | 11 | 11 | 11 |
| Family | Adjusted MS | 130.34 | 3.9849 | 516.8 | 27.83 | 30.257 | 7.4673 | 2.386 | 0.04034* | 0.1199 | 91.49 | 872.9 |
|  | DF | 72 | 72 | 72 | 72 | 72 | 72 | 65 | 63 | 72 | 72 | 64 |
| Age class | Adjusted MS | 1746.54*** | 49.2433*** | 558.3 | 11.31 | 0.756657 | 57.5553*** | 5.037* | 0.06175 | 0.1035 | 966.68*** | 594.4 |
|  | DF | 4 | 4 | 4 | 4 | 4 | 4 | 3 | 3 | 4 | 4 | 4 |
| Block | Adjusted MS | 395.44 | 20.5308* | 971.9 | 187.33* | 0.498 | 0.8034 | 3.006 | 0.18648* | 0.317910621 | 422.78* | 5954.4* |
|  | DF | 2 | 2 | 2 | 2 | 2 | 2 | 1 | 1 | 2 | 2 | 2 |
| H2 |  | 0.266 | 0.235 | 0.209 | 0.059 | 0.346 | 0.086 | 0.202 | 0.944 | -0.115 | 0.659 | 0.204 |

The narrow-sense heritability (*h^2^*) values for traits were slight-moderate with an average value of 0.281 (Table 3). The two exceptions of leaf width (0.944) and branch number (-0.115) are likely explained by the relative PEMS values for these traits, with the variation in leaf width being predominantly ascribed to differences between families and the PEMS for family in the model for branch number being negative due to family accounting for very little variation. The narrow-sense heritability for leaf width, which may already be quite high, would also be inflated by accounting for relatedness between individuals, whereas that for branch number is likely an error due to family not being an important predictor for this trait.

In the general linear models that were run for age classes one and four, region was a significant predictor in both age classes for leaf length (p=4.8 x 10^-4^ for age class one and p=2.6 x 10^-3^ for age class four). Population was a significant predictor in both age classes one and four for stem length (p=3.6 x 10^-3^ for age class one and p=3.9 x 10^-4^ for age class four). Within age class one, population was a significant predictor for spread (p=2.0 x 10^-3^). Within age class four, population was a significant predictor for stem diameter (p=7.4 x 10^-4^), number of internodes (p=7.4 x 10^-4^) and internode length (p=4.0 x 10^-3^). Family was a significant predictor for leaf width in age class four (p=3.1 x 10^-3^). Block was a significant predictor for internode length (p= 3.8 x 10^-3^) in age class one (Appendix 1).

### Correlation of trait and climate variables

In the PCA of climate variables, the first two principal components (PCs) accounted for 90.1% of the observed variation (Tables 4 and 5). PC 1 was positively associated with temperature and rainfall, especially February minimum temperature, while PC 2 was positively associated with days of ground frost and average temperature.

**Table 4:**
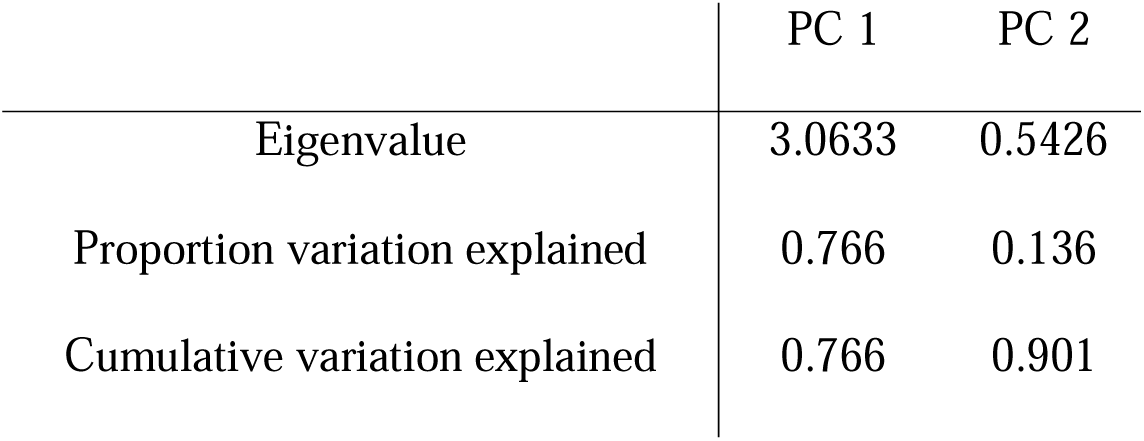
Eigenvalues and proportions of observed variation in first two principal coordinates from the PCA that was run on climate data.

|  | PC 1 | PC 2 |
| --- | --- | --- |
| Eigenvalue | 3.0633 | 0.5426 |
| Proportion variation explained | 0.766 | 0.136 |
| Cumulative variation explained | 0.766 | 0.901 |

**Table 5:** The contribution of each climate variable to the first two principal component from the PCA run on climate data.

| Variable | PC 1 | PC 2 |
| --- | --- | --- |
| Average February minimum temperature | 0.561 | 0.004 |
| Average annual cumulative rainfall | 0.489 | 0.029 |
| Average annual temperature | 0.476 | 0.684 |
| Average annual days with ground frost | -0.469 | 0.729 |

Although only the regressions against leaf width in age class one and stem diameter in age class 4 were significant, R^2^ values were as high as 79% for some of the nonsignificant regressions. R^2^ values were equal to, or larger than, 50% for stem length, stem branching, internode length and leaf width in age class one and for stem diameter, stem branching, internode length, spread and extension in age class 4 (Table 6). For each of these traits except leaf width, the coefficient was negative. PC 2 was not significant for any trait means.

**Table 6:**
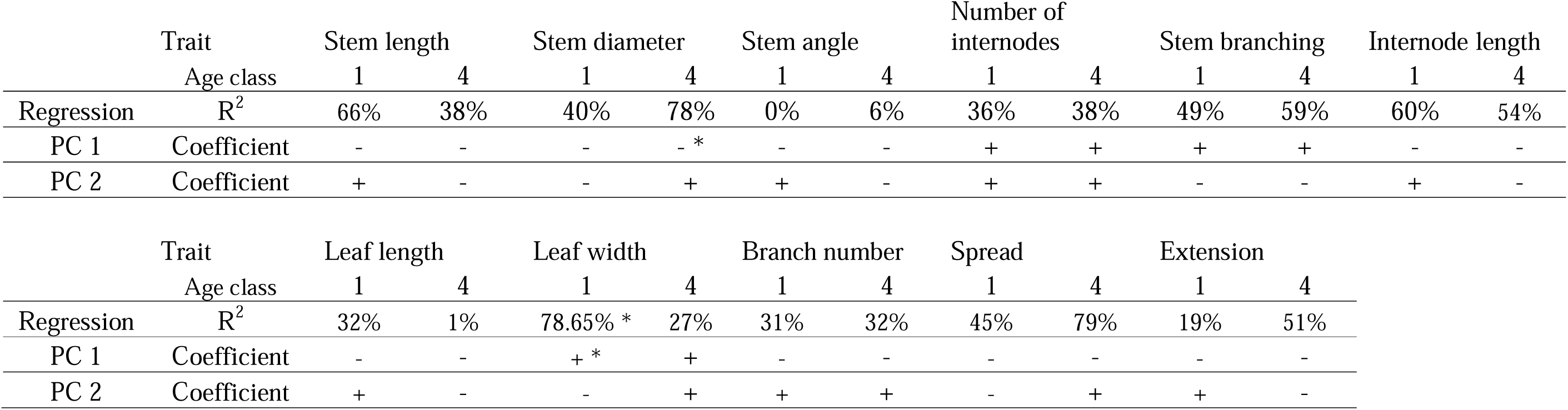
Results of principal component regression of all traits against climate principal components one and two. Significance levels for the egression and for PC 1 and PC 2 are displayed as stars next to the R^2^ value or coefficient, respectively: key: * = p-value between 0.0045 and 0.00045.

## Discussion

This paper quantifies the genetic variation in phenotypic traits of 16 natural juniper populations across juniper’s three main population centres in Britian. Our results indicate that differences between populations explains a majority of the observed variation in half of the measured traits (6 of 11), that region is an important predictor for four traits (leaf length, leaf width, branch number and extension), and that family accounts for a large proportion of variation in four traits (stem angle, leaf width, spread and extension) and is a minor component of the observed variation in all other traits. The values of narrow-sense heritability generally showed that the measured traits are slightly to moderately genetically based. Furthermore, our analyses indicate that weak but detectable correlations exist between trait variations and source population climate variables, and suggest that plants from colder, drier environments were larger than those from warmer, wetter environments when grown in the greenhouse. Finally, these correlations may be related to the age of the plants, as our comparisons between traits within the oldest and youngest age classes suggest that ontogenetic variation in resource partitioning may play a role in seedling establishment. More work is warranted to gain a full understanding of the environmental drivers of selection and how they might influence growth differently as plants age. The trial has been established outdoors in the Scottish Borders with the intention of continuing to assess these traits as the plants age.

The large differences in phenotype between populations and the similarities within them (Figure 3) indicate that British juniper populations are highly adapted to their local environments (Klimko et al., 2007; Knyazeva & Hantemirova, 2020; Rehfeldt, 1997). The observed pattern of lower among-family than among-population variation is distinctive and suggests that there have been limitations to gene flow between populations to cause their phenotypic differentiation. These findings are consistent with those of both Rehfeldt (1997) and Klimko et al. (2007), who studied the quantitative genetics of seven *Cupressus* taxa and of *J. oxycedrus* with varying levels of fragmentation, respectively, using common garden trials. The results of these studies suggest that either range contractions or genetic isolation can cause relatively higher intra-population phenotypic variation compared to inter-population variation in two species closely related to juniper. Interestingly, Scots pine and other conifers generally do not share this pattern; in Scots pine, the majority of observed variation in needle anatomy (Donnelly et al., 2016), phenology (Salmela et al., 2013) and resistance to disease (Perry et al., 2016), among other traits, were attributed to among-family variation, and among-population differences were small. Researchers have hypothesized that despite fragmentation of Scots pine populations, this is due to extensive gene flow between populations that maintains variation even in the face of strong local selection (Donnelly et al., 2016; Perry et al., 2016; Salmela et al., 2013). This contrast implies a marked difference in gene flow capability between these taxa, with juniper being relatively more limited than Scots pine.

Several caveats are worth mentioning here. Firstly, since our individuals are likely half siblings, but potentially full siblings, the phenotypic variation accounted for by families in this study is likely an underestimation of the true value by as much as half to three-quarters. Furthermore, phenotypic variation is shaped by natural selection. The pattern observed in this study, of high inter-population variation and low intra-population variation, may be due either to stabilizing selection within populations, causing a centring of phenotypes around a “more fit” mean value, or by divergent selection, where different selective pressures among populations cause phenotypic differentiation despite gene flow between them. Either or both aspects of natural selection may be responsible for increasing inter-population variation relative to intra-population variation. Alternatively, the observed pattern of phenotypic variation may be due to barriers to gene flow between populations and/or a loss of genetic diversity from inbreeding, either or both of which would also cause the observed differentiations among populations and similarities within them. Although we did not genotype the same individuals present in the common garden trial, our analysis of neutral Single Nucleotide Polymorphisms (SNPs) and Simple Sequence Repeats (SSRs) data from individuals growing in 18 natural juniper populations (Baker *et. al*, 2024; in preparation) found evidence that gene flow between populations is lower than in other northern temperate tree species, and that some genetic differentiation among populations is evident. Although stabilizing selection and different selective pressures among populations seem to be affecting phenotypic variation, the evidence of genetic differentiation and relatively limited gene flow from our analyses of neutral genetic markers suggests that genetic drift may also be a factor in the phenotypic differentiation of populations. Our genetic analysis is also in agreement with other population genetic studies of juniper across the British Isles, which found evidence for genetic differentiation of populations and barriers to gene flow (Merwe et al., 2000; Provan et al., 2008; Reynolds, 2022).

Region was a significant predictor for leaf length and width, and including regions in our models improved them for four traits: leaf length and width, number of branches and extension. For each of these traits, region accounted for approximately 20% of the observed variation with the exception of number of branches, in which region accounted for over 40% of the observed variation. Generally, plants from S. England had longer needles and those from the Lake District had wider ones. Southern English plants also displayed a distinct growth habit, with each of the nine plants having one main, upright stem (Baker et al., 2024b), which is supported by the work of Ward (2007) who studied a southern English population and found that plants grew to an average height of about 2.7 meters, with no plants shorter than 1 meter. In a previous field study, Knyazeva and Hantemirova (2020) found that needles were longer in southern Eurasian juniper trees, and used this difference as a basis for delineating the southern populations as a separate taxon. In other pine species needle morphology is an adaptively important trait in controlling a plant’s response to drought, among other things (Climent et al., 2024). Previous studies on several different pine species have consistently found a negative correlation between needle length and source habitat aridity (Grill et al., 2004; López et al., 2008; RamírezLValiente et al., 2022; Rodriguez, 2019). However, previous work on drought tolerance in junipers from across Europe found that junipers are both highly drought resistant and that populations from highly contrasting environments have largely similar hydraulic embolism resistance, suggesting that the trait is highly conserved. By contrast, previous studies in angiosperms have found that temperature and particularly growing season length, rather than aridity, is a primary driver in the adaption of leaf anatomy, with trees from warmer climates growing larger leaves (Davy & Gill, 1984; Esplugas, 2018; Grill et al., 2004; Pelham et al., 1988). The adaptation of needle anatomy in UK juniper populations therefore seems more analogous to that of angiosperms than to other conifers. This may be because juniper in the UK is closer to the northern edge of its population distribution than it is to its southern edge (Esplugas, 2018; Thomas et al., 2007), and therefore juniper needle adaptation in the UK may be driven more by colder and wetter conditions than it is by warm and dry ones. Indeed, winter precipitation has been demonstrated to be an important factor in juniper growth rate and growth habit in Europe (Carrer et al., 2019; Hallinger et al., 2010; Pellizzari et al., 2014), and may be contributing to the phenotypic adaptations across UK populations. However, whether juniper benefits from increased snowfall through increased microbial activity and insulation (Hallinger et al., 2010; Unterholzner et al., 2022), or if snowpack negatively impacts growth through shortening the growing season (Carrer et al., 2019; Pellizzari et al., 2014) is still unclear, and likely dependant on the local climate and elevation. Hallinger et al. (2010) hypothesized that these effects represent a tipping point, where some snowpack is beneficial to juniper growth, but delayed snow melting may negatively impact growth. More work is warranted on how climatic variables interact and affect the local adaptation of junipers to identify the most important factors driving adaptation in *J. communis*.

Regressions between trait variables and climate Principal Components generally lacked the statistical power to detect significant correlations. The lack of significance in these regressions is likely due to low sample sizes (N= 59 and 84 for age classes one and four, respectively) that were necessitated by having to control for age class, the fact that average trait values for each population were used to accommodate the 1km^2^ resolution of the climate data, and finally statistical overfitting of the models by including PC2. Indeed, when these PCRs were run over both age classes, all the regressions with R^2^ over 50% are significant even after the Bonferroni corrections. Although the PCRs lacked the statistical power to determine significance, the coefficients for each trait in both age classes were consistent with each other. For each of stem length, stem diameter, internode length, leaf width and spread the coefficients for PC1 between the age classes were negative and had similar values. Some trends between climate and traits were suggested in these preliminary analyses, however, they must be interpreted with caution and warrant further research. Each trait with a R^2^ value over 50% except leaf width was associated with plant growth and has a negative coefficient with both age classes, indicating that plants from colder, drier environments grew larger in the greenhouse, and that plants from warmer, wetter climates had wider leaves. This may be an effect of plants from more limiting environments requiring a lower temperature threshold to initiate growth (Salmela et al., 2013), however, in the most general terms, it demonstrates that climate is an important factor affecting the local adaptation of juniper trees.

Our work has shown a tentative correlation between climate and phenotype, suggesting local adaptation, but does not fully explore which climate factors may be the most important for the adaptation of juniper populations. Further research should be conducted to pinpoint the factors driving adaptation in juniper populations. The fact that juniper plants from more stressful environments grew larger in the common garden may be a result of relieving those stressors and/or it may be a genetic adaptation to allocate more energy to growth in harsher environments. Although we cannot distinguish whether the observed correlations between climate PCs and trait data were due relieving environmental stressors, comparisons within the oldest and youngest age classes of plants in our study hint that ontogenetic adaptation may be a factor.

The general linear models run within the oldest and youngest age classes found that populations were significantly different in multiple traits associated with growth and size within the youngest age class (stem length, stem diameter, number of internodes and internode length; p < 0.0045), but not the oldest (Appendix 1). These analyses suggest that populations were more variable in size and growth rate when they were younger, and that these differences became less pronounced as the plants aged, when they might have adopted more distinct growth habits and sizes. In the oldest age class, only stem length and spread were significantly different between populations (p < 0.0045), which may be explained by the different growth habits and sizes that the trees developed as the aged. This conclusion is consistent with other studies that hypothesize the evolution of a “go for broke” strategy where younger plants invest more in photosynthesis and transpiration to aid establishment at the expense of a higher risk of mortality (Bond, 2000; Vergeer & Kunin, 2013). More specifically, previous work evaluating age-specific growth of junipers is scarce. Kramer (1989) reported that juvenile *J. occidentalis* trees from Oregon invested more in their root systems than older individuals, and older individuals, by contrast, partitioned more of their proportional growth to their foliage. Although our study did not evaluate root growth, Kramer’s work does illustrate ontogenetic adaptation in a closely related species. Seedling life stages are likely subject to very strong selective pressure, as evidenced by the heavily skewed age distributions of natural populations towards older individuals and the reproductive failure of natural populations (Thomas et al., 2007). These differences between age classes in our data hint at the possibility that energy partitioning to growth may be both adaptive and age-specific. More detailed work is needed to understand the ecological trade-offs that may influence this putative adaptive life history pattern.

### Conservation recommendations

Conservation managers face a difficult situation in protecting the remnant juniper stands in the UK, particularly with the introduction of the novel pathogen *Phytophthora austrocedri* (Green et al., 2015). Although there is some evidence for natural resistance to *P. austrocedri* among UK juniper populations (Green et al., 2020), the capacity for natural populations to develop this resistance may be hampered by the lack of or low levels of natural regeneration and genetic isolation of populations. Furthermore, planting operations may play a role in the spread of *P. austrocedri* to populations within 2km of each other (Donald et al., 2021) or even inadvertently damage the established juniper trees (De Frenne et al., 2020). Planting into existing populations is therefore strongly discouraged. Encouraging natural regeneration in established populations, for example using fencing to exclude grazers or sod cutting to create disturbance (De Frenne et al., 2020; Verheyen et al., 2005) is likely the best way to facilitate the adaptive potential of juniper while mitigating the risk of *P. austrocedri*.

Although the pollen and seed dispersal of juniper is not fully understood, our work has demonstrated that juniper probably has less effective gene flow than other conifer species, which is evident in the lower levels of within-population compared to between-population variation, and which is also supported by our parallel work on neutral genetic markers (Baker *et al*. 2024; in preparation). Therefore, the establishment of “satellite” populations – small, planted populations interspersed among remnant fragments at a minimum distance of 1km from existing populations – may facilitate gene flow, reduce the effects of inbreeding and help to maintain the adaptive potential of populations. This paper has also illustrated the importance of local adaptations to junipers, evidenced by the differences in adaptive phenotypes among populations, which have been selected for by their respective environments. In light of this, potential satellite populations should be composed of locally sourced material to reduce the chances of environmental mismatch. Stock for satellite populations should also be raised from seeds, rather than cuttings, to reduce the effects of inbreeding. Future work should focus on juniper’s gene dispersal capacity and reconnecting population fragments. Although the guidance to source & grow locally may appear to contradict a call to increase connectivity, we recommend that wherever possible natural gene flow connections are encouraged, rather than transplantations, to minimise the biosecurity risks associated with moving plant materials. Finally, this paper shows the distinct phenotypic differentiation of juniper populations across Britain. The authors recommend that several of the populations across the range studied here are considered for designation as Gene Conservation Units under the EUFORGEN framework (Hubert & Cottrell, 2014) based on both the findings presented here and in our companion paper on neutral genetic markers (Baker *et al*. 2024; in preparation).

## Data accessibility statement

The data described in this article are accessible open-access through the Environmental Information Data Centre (EIDC). Please see Baker et al. (2024b) for the trait data described here, and (Baker et al., 2024a) for the germination and seedlot collection data for the full trial.

## Competing interests statement

The authors declare no conflicts of interest.

## Author contributions

**James Baker:** Data curation (lead), formal analysis (lead), visualization (lead), writing-original draft (lead), writing-review and editing (equal). **Joan Cottrell:** Conceptualization (equal), project administration (equal), writing-review and editing (equal), supervision (equal). **Richard Ennos:** Conceptualization (equal), project administration (equal), writing-review and editing (equal), supervision (equal). **Annika Perry:** Conceptualization (equal), project administration (equal). **Sara Green:** Sample collection (lead), project administration (equal). **Stephen Cavers:** Conceptualization (lead), project administration (lead), writing-review and editing (equal), supervision (lead).

### Acknowledgements

The authors are very grateful to the funders of this project, The Scottish Forestry Trust, Forest Research, The Botanist Foundation, The Woodland Trust and the UKCEH. We are also very thankful for the hard work of Elanor James, who was responsible for gathering phenotypic data for the trial.

## Notes

https://catalogue.ceh.ac.uk/documents/330cf3ac-21c3-4fa8-ab76-528f8cb2fbb8

https://catalogue.ceh.ac.uk/documents/f61dcdcc-b838-4bfa-8d94-7820114a68c8

### Competing Interest Statement

The authors have declared no competing interest.

