## Appendix for "Not like other conifers: evaluation of phenotypic diversity in British common juniper, *Juniperus communis*, indicates genetic isolation and local adaptations among remnant populations"

Appendix 1: Results from general linear models run for all traits within the youngest (1) and oldest age (4) classes. Tests were nested general linear models using regions (fixed factor), populations (fixed factor) nested within regions, families (random factor) nested within populations and block as a random control factor. Stars next to adjusted MS values indicate significance: key: * = p-value between 0.0045 and 0.00045, ** = p-value between 0.00045 and 0.000045

|  | Trait | Stem length | | Stem diameter | | Stem angle | | Number of internodes | | Stem branching | | Internode length | |
| --- | --- | --- | --- | --- | --- | --- | --- | --- | --- | --- | --- | --- | --- |
|  | Age class | 1 | 5 | 1 | 5 | 1 | 5 | 1 | 5 | 1 | 5 | 1 | 5 |
|  | Model Adjusted R^2^ | 31.4% | 30.6% | 19.9% | 20.2% | 7.6% | 10.3% | 0.0% | 26.5% | 25.9% | 9.1% | 33.8% | 16.6% |
| Region | Adjusted MS | 101.37 | 123.4 | 0.545 | 0.486 | 612.9 | 795.8 | 10.97 | 17.62 | 31.207 | 43.641 | 6.295 | 19.23 |
|  | DF | 2 | 2 | 2 | 2 | 2 | 2 | 2 | 2 | 2 | 2 | 2 | 2 |
| Population | Adjusted MS | 355 * | 604.8 ** | 8.03 | 13.1738 * | 796.1 | 610.1 | 23.84 | 112.22 * | 56.742 | 47.606 | 10.781 | 26.933 * |
|  | DF | 12 | 9 | 13 | 9 | 12 | 9 | 12 | 9 | 12 | 9 | 12 | 9 |
| Family | Adjusted MS | 122.17 | 143.9 | 3.759 | 3.327 | 415.3 | 540.7 | 22.42 | 28.83 | 27.578 | 25.559 | 10.657 | 8.432 |
|  | DF | 41 | 40 | 41 | 40 | 41 | 40 | 41 | 40 | 41 | 40 | 41 | 40 |
| Block | Adjusted MS | 81.17 | 133.5 | 4.743 | 6.256 | 251.1 | 497.3 | 67.79 | 62.69 | 8.525 | 1.546 | 34.843 * | 1.804 |
|  | DF | 2 | 2 | 2 | 2 | 2 | 2 | 2 | 2 | 2 | 2 | 2 | 2 |
|  | Trait | Leaf length | | Leaf width | | Branch number | | Spread | | Extension | |  |  |
|  | Age class | 1 | 5 | 1 | 5 | 1 | 5 | 1 | 5 | 1 | 5 |  |  |
|  | Model Adjusted R^2^ | 47.1% | 35.2% | 50.0% | 53.7% | 21.8% | 2.6% | 32.4% | 18.3% | 12.0% | 27.3% |  |  |
| Region | Adjusted MS | 20.071 * | 16.1453 * | 0.206 | 0.247 | 0.447 | 0.117 | 59.54 | 174.16 | 375.9 | 228.9 |  |  |
|  | DF | 2 | 2 | 2 | 2 | 2 | 2 | 2 | 2 | 2 | 2 |  |  |
| Population | Adjusted MS | 4.792 | 6.0873 | 0.0440 | 0.0322 | 0.242 | 0.253 | 215.76 * | 131.08 | 588.5 | 769 |  |  |
|  | DF | 12 | 9 | 12 | 9 | 12 | 9 | 12 | 9 | 12 | 9 |  |  |
| Family | Adjusted MS | 2.34 | 2.5361 | 0.0338 | 0.0507 * | 0.132 | 0.093 | 67.25 | 67.68 | 828 | 927.7 |  |  |
|  | DF | 33 | 37 | 33 | 37 | 41 | 40 | 41 | 40 | 38 | 39 |  |  |
| Block | Adjusted MS | 2.658 | 0.0025 | 0.0912 | 0.00403 | 0.217 | 0.224 | 152.56 | 179.24 | 3495 | 1821.6 |  |  |
|  | DF | 1 | 1 | 1 | 1 | 2 | 2 | 2 | 2 | 2 | 2 |  |  |

Appendix 2: Climate values for each population used in the PCR with the overall average and standard deviation (Stdev).

| Pop | Febuary min. temp (°C) | Annual cumulative rainfall (mm) | Average annual temp (°C) | Annual cumulative days with groundfrost (days) |
| --- | --- | --- | --- | --- |
| AG | 1.93 | 2017.36 | 8.62 | 82.46 |
| AR | 1.71 | 2126.64 | 8.69 | 111.97 |
| BG | 0.30 | 956.14 | 8.59 | 138.50 |
| GA | 0.49 | 1407.27 | 8.68 | 132.54 |
| IN | 1.02 | 1113.89 | 8.13 | 89.06 |
| LM | 0.10 | 1051.33 | 7.32 | 124.54 |
| MB | -1.53 | 1059.20 | 6.84 | 144.73 |
| TY | 0.84 | 1394.97 | 8.30 | 118.63 |
| WW | -0.55 | 880.66 | 7.46 | 139.53 |
| Average | 0.72 | 1473.49 | 8.46 | 116.86 |
| Stdev | 0.9486 | 658.76 | 0.9959962 | 20.206172 |
